# BD5: an open HDF5-based data format to represent quantitative biological dynamics data

**DOI:** 10.1101/2020.04.26.062976

**Authors:** Koji Kyoda, Kenneth H. L. Ho, Yukako Tohsato, Hiroya Itoga, Shuichi Onami

## Abstract

BD5 is a new binary data format based on HDF5 (hierarchical data format version 5). It can be used for representing quantitative biological dynamics data obtained from bioimage informatics techniques and mechanobiological simulations. Biological Dynamics Markup Language (BDML) is an XML(Extensible Markup Language)-based open format that is also used to represent such data; however, it becomes difficult to access quantitative data in BDML files when the file size is large because parsing XML- based files requires large computational resources to first read the whole file sequentially into computer memory. BD5 enables fast random (i.e., direct) access to quantitative data on disk without parsing the entire file. Therefore, it allows practical reuse of data for understanding biological mechanisms underlying the dynamics.

## Introduction

Recent advances in bioimage informatics and mechanobiological simulation techniques have led to the production of a large amount of quantitative data of spatiotemporal dynamics of biological objects ranging from molecules to organisms [1]. A wide variety of such data can be described in an open unified data format Biological Dynamics Markup Language (BDML), an Extensible Markup Language (XML)-based format [2]. BDML enables efficient development and evaluation of software tools for a wide range of applications.

The XML-based BDML format has the advantages of machine/human readability, and extensibility. However, it is often problematic for accessing and retrieving data when the size of the BDML file becomes too large (e.g., our programs cannot load a BDML file over 20 GB on a standard workstation). This problem arises because parsing an XML-based file often requires large computational resources to first read the whole file sequentially into computer memory. In fact, many sets of quantitative data stored in the SSBD:database (Systems Science of Biological Dynamics database) [1] were divided into a series of BDML files for each time point to allow software to read them efficiently. One of the solutions to the above problem is to use another approach such as the eXtensible Data Model and Format [3] or FieldML [4]. In these formats, the data itself is described in HDF5 binary format and meta-information about the data is described in XML format. HDF5 is a hierarchical data format for storing large scientific data sets (http://www.hdfgroup.org/HDF5/). It is widely used for describing various kinds of large-scale biological data [4-9].

Here, we describe the development of BD5 data format, based on HDF5, for representing quantitative biological dynamics data in a manner that enables quick access and retrieval.

## Materials and Methods

### Design and implementation

Here, we extended BDML to support HDF5-based storage of quantitative biological dynamics data. In contrast to XML documents, HDF5 format can allow random (i.e., direct) access to parts of the file without parsing the entire contents. Therefore, HDF5 is a more efficient file format for accessing and retrieving the contents of the file.

We developed the BD5 data format based on HDF5 for representing quantitative data. A BD5 file is organized into two primary structures, *datasets* and *groups*. Datasets are array-like objects that store numerical data, whereas groups are hierarchical containers that store datasets and other groups. Detailed information on BD5 is available at http://ssbd.qbic.riken.jp/bdml/. Here, we summarize the BD5 major datasets and groups. BD5 format has one container named *data* (Fig. 1). It includes

- *scaleUnit* dataset for the definition of spatial and time scales and units,
- *objectDef* dataset for the definition of biological objects,
- *featureDef* dataset for features of interest,
- numbered groups (0, 1, …, n) corresponding to an index number of a time-ordered sequence,
- *trackInfo* dataset for the information of tracking of one object to another.

**Figure 1.**
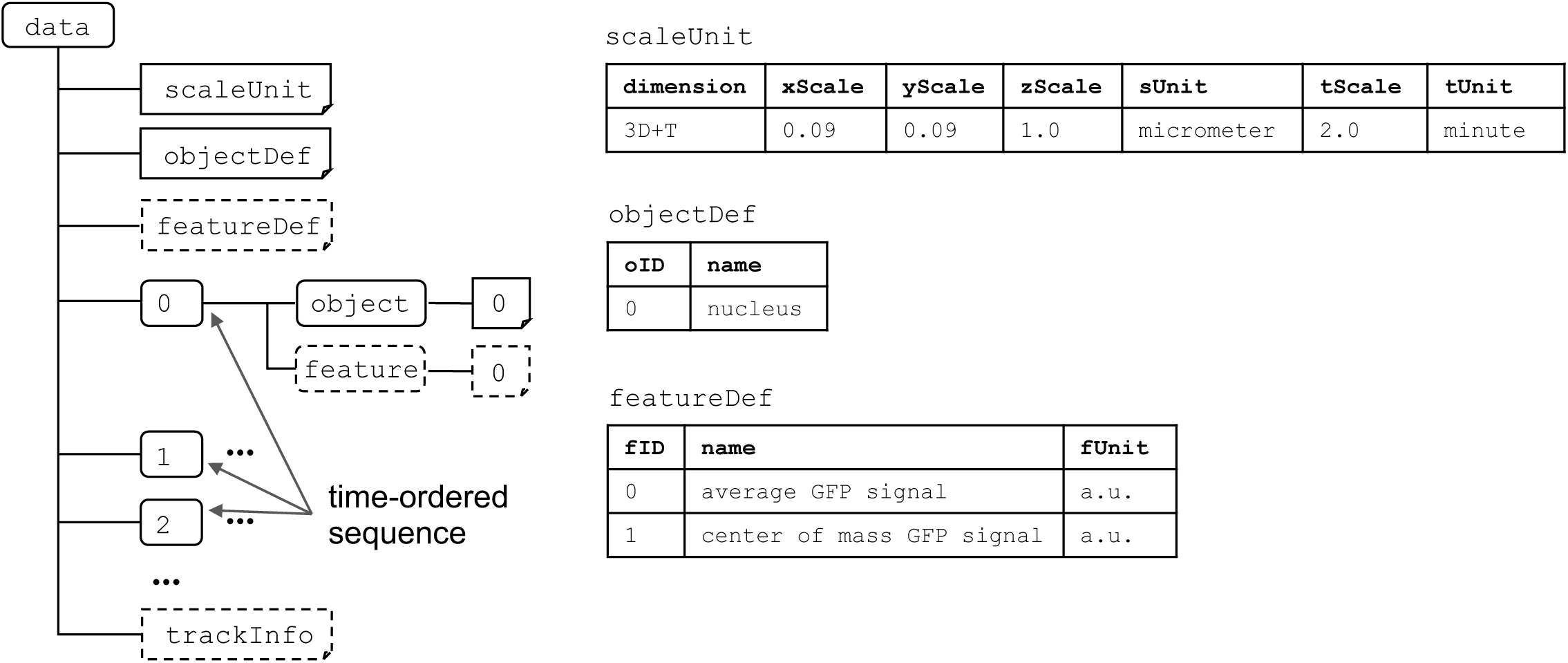
Outline of the BD5 data format. The *data* group includes *scaleUnit, objectDef, featureDef*, and *trackInfo* datasets; each *data* group is numbered to correspond to the index number of the time-ordered sequence. Each numbered group has spatial information about biological objects and numerical information about features related to the objects. Solid and dashed boxes represent the required and optional elements, respectively.

Each of the numbered groups corresponds to an index of a time-ordered sequence that has *object* and *feature* groups. For a fixed time interval, the index will correspond to each sequential time point. For example, if the time interval is 2 minute, group 0 will have t = 0 and group 1 will have t = 1 while tScale is 2 and tUnit is minute (Fig. 1). For irregular time intervals, the index allows a time-ordered sequence to be saved and be read in the correct order. If the first time is 0 minutes, the second time is 2 minutes, while the third time is 7 minutes, then group 0 will have t = 0, group 1 will have t = 2 and group 2 will have t = 7. The tUnit is still minute, but the tScale in this case will be 1.

Each *object* group has numbered dataset(s) corresponding to the reference number of the biological object(s) predefined under the *objectDef* dataset. Each row of the numbered object includes an identifier of the object and its spatiotemporal information such as time point and *xyz*-coordinates (Fig. 2). To represent biological objects such as line and face entities that have an arbitrary number of *xyz*-coordinates in BD5 format, a tabular dataset is used (Fig. 3). The multiple *xyz*-coordinates are represented by using a sequential ID (sID) that allows us to connect the *xyz*-coordinates together to form a line or a face within a biological object.

**Figure 2.**
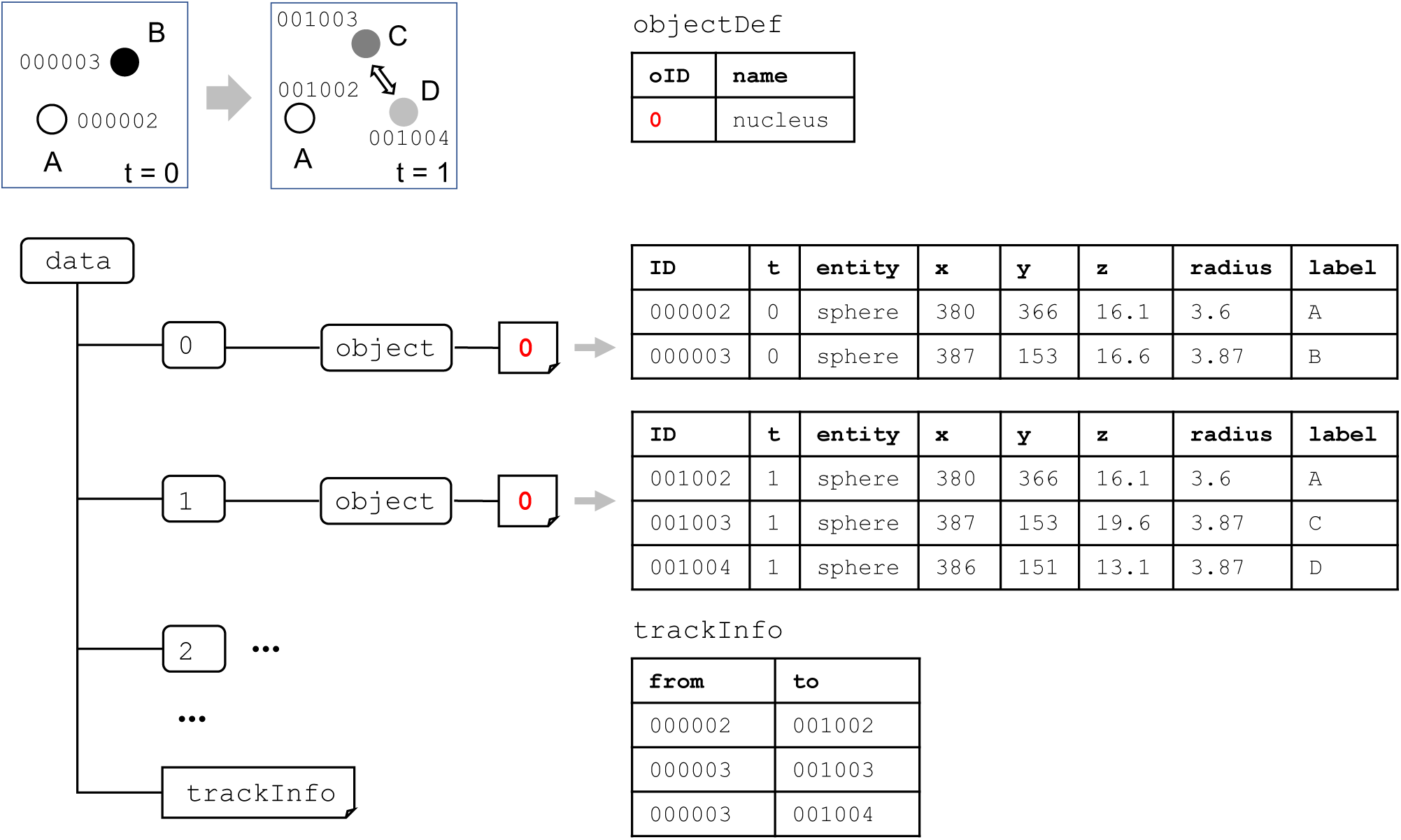
An example of the description of the spatiotemporal information of biological objects and their tracking information. The dataset name in *object* group corresponds to the identifier (ID) of the biological object (red). Each row in the dataset must have a unique ID and its spatiotemporal information. A label can optionally be attached for each object. The tracking information including object divisions and fusions can be stored in *trackInfo* dataset.

**Figure 3.**
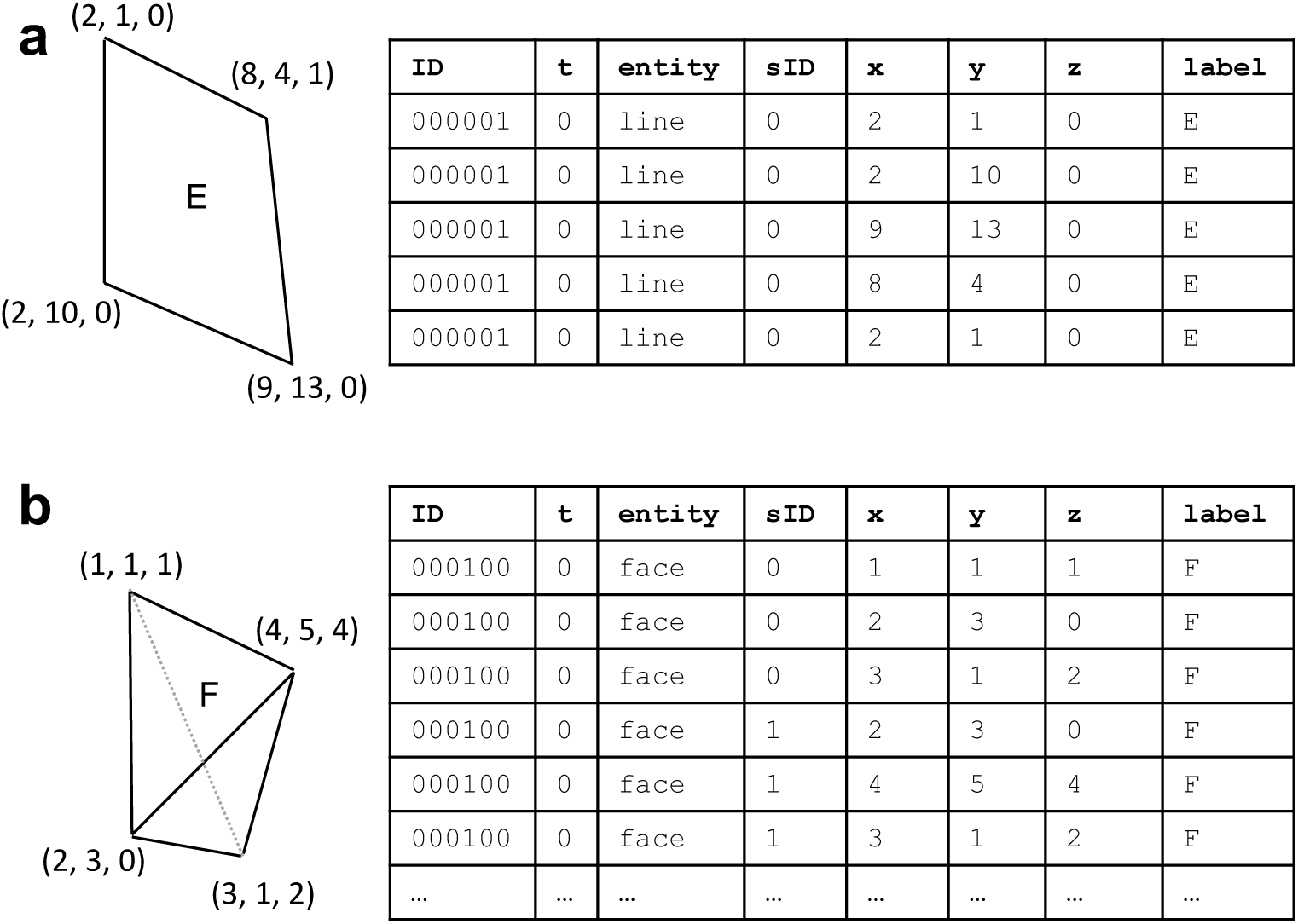
An example of the description of the spatiotemporal information based on line (a) and face entities (b). The sequential identifier (sID) represents a set of coordinates that can be connected beginning at the top to describe an entity within one biological object.

Each *feature* group has numbered dataset(s) corresponding to the reference number of the object(s) predefined in the *objectDef* dataset. Each row of the numbered object includes an identifier of the object, an identifier of the feature (fID) predefined in *featureDef*, and the value of the feature (Fig. 4). This format allows objects that do not possess all the features defined in *featureDef* to be recorded, because not all the features can necessarily be measured in practical biological experiments. For example, in the experiment in Fig. 4, an object may have information for fID = 1 (name: center-of-mass GFP signal) but not fID = 0 (name: average GFP signal).

**Figure 4.**
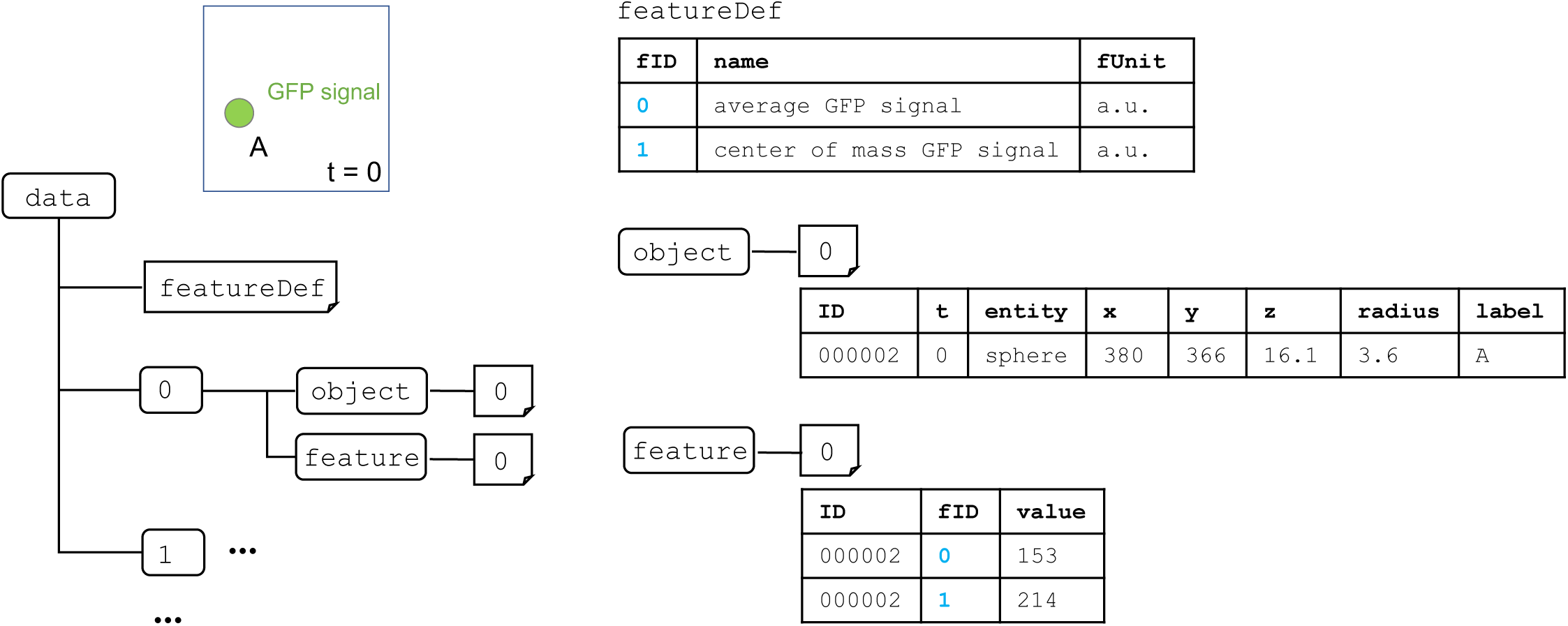
An example of the description of the feature information related to biological objects. The example object is a nucleus expressing green fluorescent protein (GFP) at t = 0 in a time series. The dataset name in *feature* group corresponds to the identifier (ID) of the biological object. Each row in the dataset has object ID, feature fID (blue), and the feature value. In this example, fID is 0 or 1 depending on whether the data is total or average GFP signal, respectively. Object ID is 0 if the object is a nucleus. Feature value is the fluorescence intensity expressed in a.u. (arbitrary units).

The *trackInfo* dataset enables information of the objects to be linked between different time points or time frames (Fig. 2). For example, when a cell at t = 0 divides into two daughter cells at t = 1, it has links from the parent cell to the daughter cells. The *trackInfo* dataset can be used to represent not only phenomena such as cell division but also those such as cell fusion.

To allow the use of BD5 to describe quantitative data, we needed to update the BDML format so that it could be used to describe the corresponding meta-information. The latest version of BDML (version 3.0) can handle an external file by using the `extFile` element (Fig. 5). The `bd5File` element that we introduced within the `extFile` element can be used to point to an external BD5 file. In addition, this update allows the designation of multiple contact persons and the use of a unique persistent digital identifier, ORCID (https://orcid.org) in its format.

**Figure 5.**
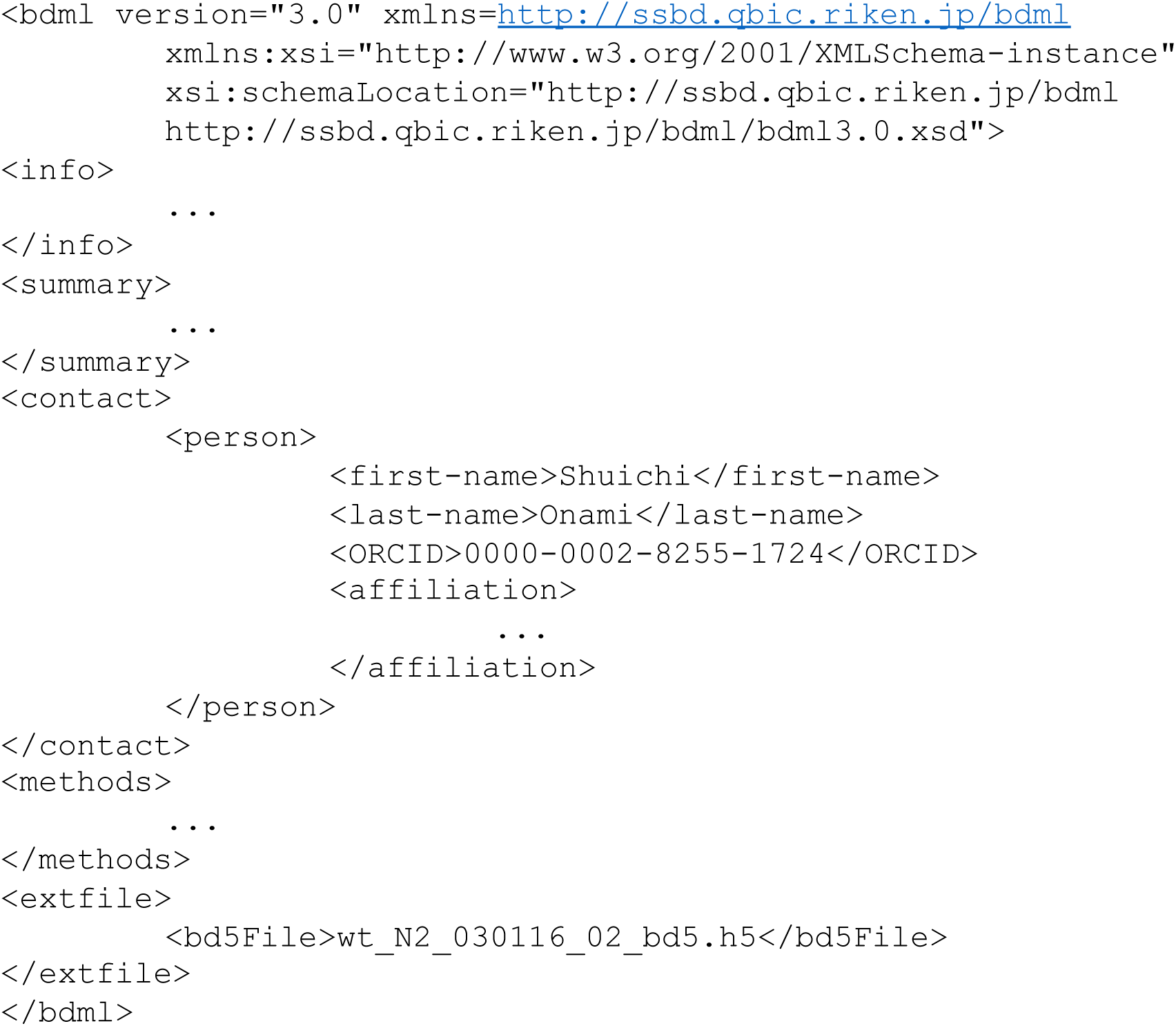
A skeleton of a BDML version 3.0 file for describing meta-information. This version allows the use of an external file for describing the data itself, designation of multiple contact persons, and the use of ORCID, a unique persistent digital identifier of the research scientist.

## Results

### Validation

To evaluate the performance of the BD5 format, we first compared time for accessing the file between XML- and HDF5-based files (i.e., between pairs of BDML and BD5 files containing equivalent data). We measured the time for accessing coordinate data at a randomly selected time point in the BDML and BD5 files (334 pairs of files) by using a Python-based program (Fig. 6a). The results indicate that the access times of HDF5- based files were consistently faster than those of the corresponding XML-based files. Therefore, BD5, the new HDF5-based format, enables practical access to quantitative data for further analysis.

**Figure 6.**
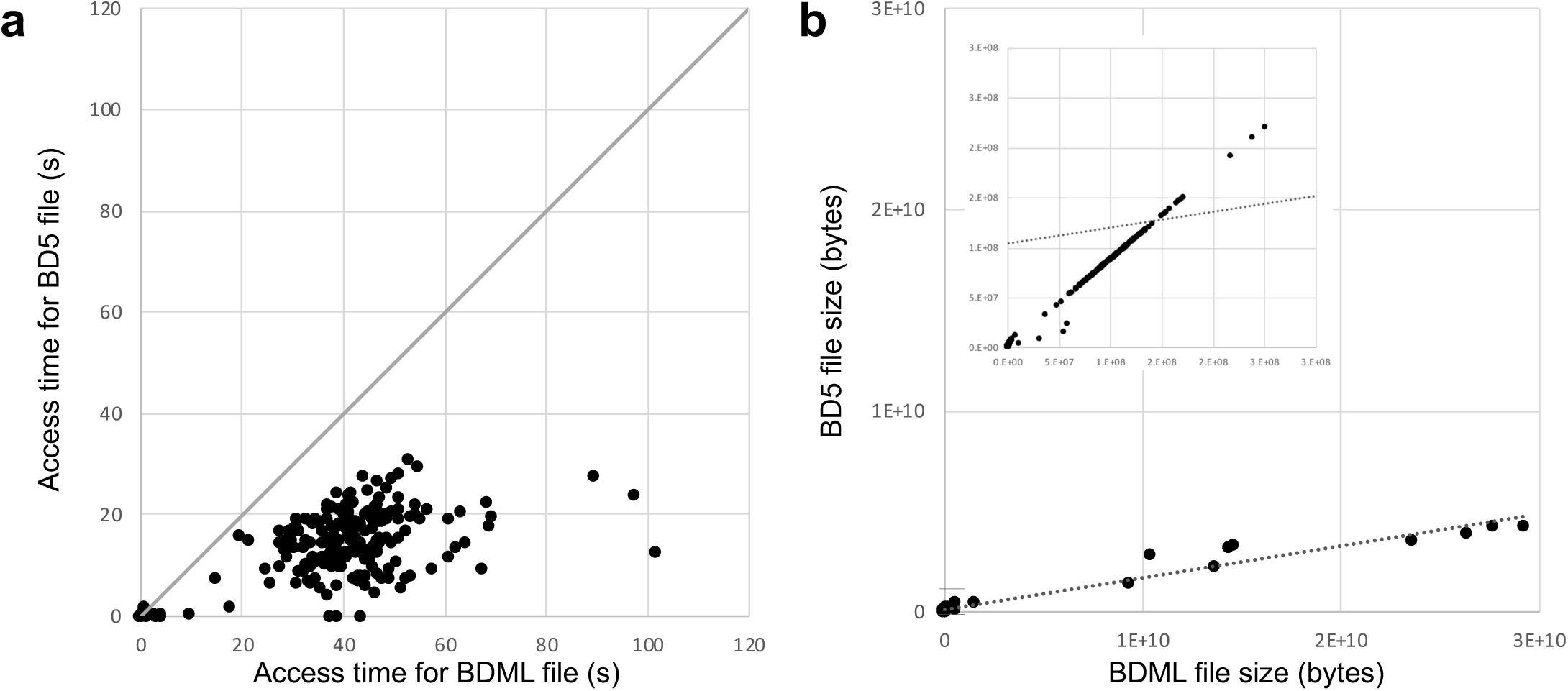
Comparison between BD5 and BDML data formats. a) Access times of the BDML and BD5 files. Access time was measured as the time for accessing and displaying *xyz*-coordinate data at a randomly selected time point stored in the BDML and BD5 files. The time was measured on an Intel Xeon CPU 2.8 GHz processor with 32 GB of main memory. Each dot represents a biological quantitative data set. We used 334 biological quantitative data sets, each of which has coordinate data and is stored in SSBD:database as a single BDML file. BD5 files were generated from the BDML files by using the BDML2BD5 program. b) Size of the BDML and BD5 files. Each dot represents a biological quantitative data set. In this comparison, we used 450 biological quantitative data sets stored in SSBD:database as BDML files. As above, the BD5 files were generated from the BDML files by using the BDML2BD5 program. The dashed line represents the linear regression line for all dots. The data within the small rectangle near the origin in the large graph is plotted on expanded axes in the insert.

File size can be a critical benchmark for a data format because the transfer of large files often fails. Therefore, we next compared disk space requirement between the XML- and HDF5-based files by comparing the size of BDML and BD5 files (450 pairs of files) (Fig. 6b). BD5 format reduced the file size by ∼85% compared with the BDML format when the data is large. When the data is small (< 300MB), the size of BD5 file is close to, but still less than, that of the corresponding BDML file. Because the size of HDF5-based files for large data is much less than that of the equivalent XML-based files, the BD5 format enables, in theory, fast transfer of large quantitative data to and from computers on the network and on the internet.

In addition, we determined the relationships between access time and file size for BDML and BD5 files (Fig. 7). In BD5, we found fast access to the coordinate data even when the file size was large. This fast data access in BD5 originated from its random access to data. In BDML, the access time linearly increased with file size. This result suggests that parsing of XML was the main bottleneck of data access. Quantitative biological dynamics data tends to be large due to the advances in live-cell imaging techniques and imaging equipment. We anticipate that BD5 will play a key role in fast access to such large data sets.

**Figure 7.**
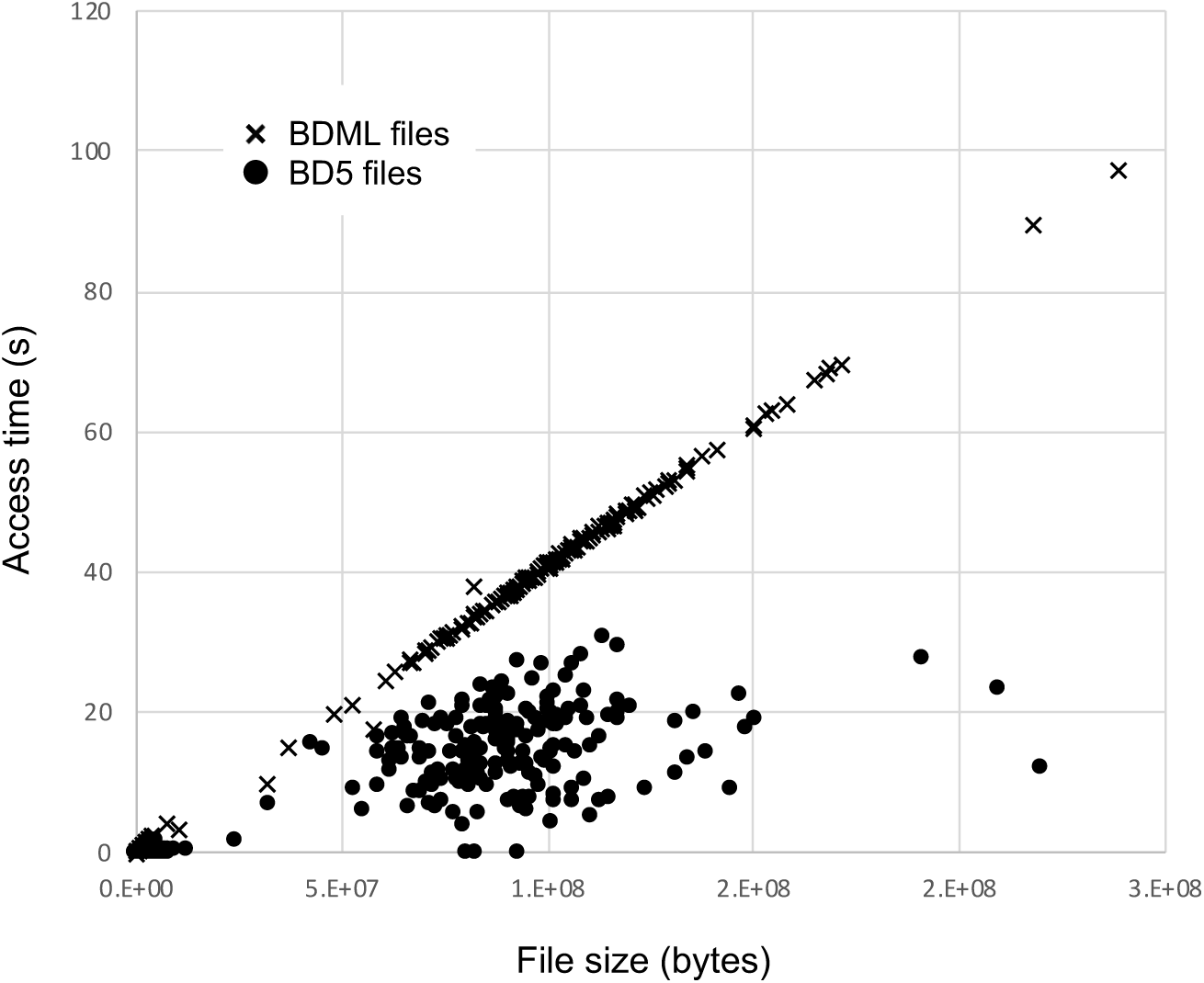
Relationship between access time and file size for BDML and BD5 files. The time for accessing and displaying coordinate data at a randomly selected time point is plotted against file size. Each cross represents a BDML file; each dot represents a BD5 file. In the comparison, we used BDML and BD5 files of the 334 biological quantitative data sets described in Fig. 6a.

### Software tools and usage related to BD5

So that BD5-based tools can be used for data stored in older BDML files, we provide a C++-based software tool named BDML2BD5. By using this tool, BDML files can be converted into BD5 files. To compile the tool, the HDF5 library is required for HDF5 data writing, and CodeSynthesis XSD (http://www.codesynthesis.com/products/xsd/) is required for the BDML schema to C++ data binding compiler. All source codes and the executable file of BDML2BD5 are available at https://github.com/openssbd/BDML2BD5/.

We also provide a program bd5lint for detecting bugs and inconsistencies in BD5 files. The program checks the structure of BD5 files, and checks that the ordered numbered datasets in *object* and *feature* groups correspond to the reference numbers of the objects and features predefined in the *objectDef* and *featureDef* dataset. It also checks the consistency of the dimensions declared and the actual dimensions used within the datasets. It provides type checking of the data and error warnings if the data do not conform to the BD5 specification. The Python source code is available at https://github.com/openssbd/bd5lint/.

We also provide several Python-based programs for data analysis using BD5 files. These programs are available as Jupyter Notebook files at https://github.com/openssbd/BDML-BD5/. An example is a program that counts the number of biological objects in each numbered group of the time-ordered sequence. By using this program, we can obtain the proliferation curve of *Caenorhabditis elegans* embryogenesis. The program can be modified to obtain similar information for other organisms such as *Danio rerio* and *Drosophila melanogaster*.

## Discussion

In this study, we developed a new BD5 data format based on HDF5 for representing quantitative biological dynamics data. Compared with BDML, which is based on XML, the BD5 format has two advantages: (a) fast access and retrieval to quantitative data because of random access to the HDF5-based file, and (b) fast transfer of files containing large quantitative data because the file size is dramatically reduced. A drawback of the BD5 is that human readability is low when compared with BDML format. BD5 files cannot be opened by text editors because the file is binary formatted. However, the HDF group provides a software tool named HDFView that enables the user to open and read all HDF5-based files (https://www.hdfgroup.org/downloads/hdfview/). This tool can compensate for the lack of human readability.

BD5 format has already been used in the latest version of SSBD:database (http://ssbd.qbic.riken.jp), which is one of the major databases for sharing bioimage data and quantitative biological dynamics data [10]. Over 687 files, which include a wide variety of quantitative biological dynamics data from molecules to cells to organisms, are available. This demonstrates that the BD5 format has high functionality and flexibility for representing quantitative biological dynamics data. SSBD:database also provides a RESTful API (i.e., an API (application programming interface) that allows applications to access data and interact with external software tools) through the use of the webservice h5serv (https://github.com/HDFGroup/h5serv). This enables SSBD:database to provide a web service for users to access quantitative data stored in BD5 files (http://ssbd.qbic.riken.jp/restfulapi/). Because HDF5 and XML are supported by many software platforms, BD5 is a promising data format for storing quantitative biological dynamics data.

Like BDML, BD5 can represent quantitative biological dynamics data that is associated with, but independent of, microscopy images. Such data has often been represented as regions of interest (ROIs) on the corresponding microscopy images; for example, the ROIs in the OME data model (https://docs.openmicroscopy.org/ome-model/) and segmentation channels in Cell Feature Explorer (https://cfe.allencell.org). However, not all data can be represented as an ROI on a microscopy image. For example, in an automated cell lineage tracing study of *Caenorhabditis elegans*, each nucleus was represented as a sphere with center and radius, independently of the z-stack images [11]. Such flexible representation of BD5 (and also BDML) enables us to represent quantitative biological dynamics data obtained not only from bioimage informatics but also from mechanobiological simulation techniques.

## Acknowledgements

We are grateful to the members of the Onami laboratory, RIKEN Center for Biosystems Dynamics Research, Japan for feedback and discussions.

